# Societal attention toward extinction threats

**DOI:** 10.1101/706911

**Authors:** Ivan Jarić, Céline Bellard, Franck Courchamp, Gregor Kalinkat, Yves Meinard, David L. Roberts, Ricardo A. Correia

## Abstract

Public attention and interest in the fate of endangered species is a crucial prerequisite for effective conservation programs. Societal awareness and values will largely determine whether conservation initiatives receive necessary support and lead to adequate policy change. Using text data mining, we assessed general public attention toward climate change and biological invasions, which represent two key threats for endangered amphibian, reptile, bird and mammal species in three European countries (France, Germany and the United Kingdom). Our analysis revealed that public attention patterns differed among species groups and countries but was globally higher for climate change than for biological invasions. Both threats received better recognition in threatened than in non-threatened species, as well as in native species than in species from other countries and regions. The relative level of attention toward the two threats corresponded well with the listing of the threats and their severity within the IUCN Red List. We conclude that more efficient communication regarding the threat from biological invasions should be developed, and that conservation practitioners should take advantage of the existing attention toward the threat of climate change.

## 1. Introduction

With an ongoing global biodiversity crisis, it is critical to overcome the present gap between research and implementation in conservation (Toomey et al. 2017). Failure to halt biodiversity decline does not solely come from a lack of scientific understanding of drivers of decline, but also from a lack of societal support and action (Schultz, 2011). Societal awareness and values partly determine the level of support for, engagement with and effectiveness of conservation initiatives, as the societies tend to protect only what they recognize as important (Stokes, 2007; Kim et al., 2014).

The emerging field of conservation culturomics provides unprecedented opportunities to understand societal attention and interests related to conservation objectives (Ladle et al., 2016; Sutherland et al., 2018). Culturomics aims to generate new insights in human culture through the quantitative analysis of digital data. Conservation culturomics is focused on the understanding of cultural dynamics associated with conservation, and comprises diverse sources and approaches, including webpage retrieval analysis (Internet salience), assessment of web search activity, page views of digital archives such as Wikipedia, contents of social networks and online news, and images and videos posted online (Roll et al., 2016; Correia et al., 2018; Jarić et al., 2019a; Retka et al., 2019; Toivonen et al., 2019). A number of studies within this field have recently focused on public attention and interest in major anthropogenic impacts on natural systems, such as climate change and invasive species, based either on web search query data or online news media (McCallum and Bury, 2013; Anderegg and Goldsmith, 2014; Funk and Rusowsky, 2014; Proulx et al., 2014; Veríssimo et al., 2014; Nghiem et al., 2016; Burivalova et al., 2018; Legagneux et al., 2018; Correia et al., 2019; Troumbis, 2019). However, as opposed to the intensive research of general societal interest in these threats, there were no attempts so far to assess societal attention toward extinction threats related to specific species and species groups. Such studies can provide valuable insights regarding public recognition of the actual threats, and major attention gaps and biases. Furthermore, higher attention toward extinction threats faced by a particular species is likely to also contribute to perceived relevance of the species for the society and its overall visibility.

The aim of this study was to explore how much attention is directed at climate change and biological invasions in relation to different species groups. We focus on climate change and biological invasions as two key processes that affect biological diversity and are often interrelated and act synergistically (Dukes and Mooney, 1999; Clavero and Garcia-Berthou, 2005; Walther et al., 2009; Bellard et al., 2012, 2018; McClelland et al., 2018). We hypothesize that, because biological invasions are not yet very well understood, they generate less societal attention than climate change, which can be a basic hindrance for conservation (Courchamp et al., 2017). We test this by comparing the Internet salience of biological invasions with that of climate change, which is expected to be characterized by a much higher level of public attention. Based on webpage retrieval analysis, we assessed the Internet salience of these two threats in relation to threatened amphibian, reptile, bird and mammal species in France, Germany and United Kingdom. Our results provide novel insights into the differences in the societal attention directed at these two threats, among different countries, species and their origin, as well as in the level of attention overlap with the actual existence and severity of these threats. We address the implications of the results for conservation management and outreach planning.

## 2. Methods

Our analysis was based on the Internet salience (i.e. the volume of Internet pages containing a particular term; Jarić et al., 2016; Correia et al., 2017) of terms referring to extinction threats in relation to multiple species. Internet salience is considered a good indicator of a species’ cultural visibility (Correia et al., 2016; 2017), and it is therefore likely that this capacity also applies to species extinction threats. Furthermore, the Internet is increasingly used as a source of information (Miller, 2005; Davies et al., 2018), and because most Internet content is generated by self-interest (e.g. blogs, Wikipedia, social media) or perceived interest (e.g. news outlets), more available content is likely to generate more reads and thus more attention. Therefore, there is a likely strong feedback loop between content availability and public interest and attention (Ladle et al., 2016).

We compiled a dataset with species scientific names (including synonyms), taxonomy, Red List categories, threat classification and threat severity (i.e., the overall decline caused by the threat) based on data from the IUCN Red List database (IUCN, 2017). The dataset comprised 5988 threatened mammal, bird, reptile and amphibian species (i.e. classified as VU, EN or CR). National red lists of Germany, France and United Kingdom, including both regionally threatened and non-threatened species from the four taxon groups (excluding species from overseas territories), were also downloaded from the National Red List database (http://www.nationalredlist.org - downloaded on 14 December 2017). Threatened species in Germany were considered to be those with the regional status being either “Declining Species”, “Possibly Endangered”, “Endangered”, “Critically Endangered”, “Extremely Rare”, or “Threatened with Extinction”; in United Kingdom, it was either “Amber” or “Red” regional status, and “Vulnerable”, “Endangered” or “Critically Endangered” in France. Since one of the objects of the study was to compare Internet salience of threats for threatened and non-threatened species from each country, and the German national red list comprised exclusively threatened species, a list of non-threatened species from Germany was obtained from the global IUCN Red List database (IUCN, 2017).

Internet salience data collection was performed during March 2018, in line with the approach proposed by Jarić et al. (2016) and Correia et al. (2017). Search was conducted using scientific species names, as they represent a reliable proxy and preferable alternative to vernacular names, due to a strong and culturally independent association between their representation in digital corpora (Jarić et al., 2016; Correia et al., 2017). Google’s Custom Search Engine API was used to search websites from each of the three countries for pages mentioning scientific species names, terms representing the two threats (i.e., climate change and invasive species, respectively), and their combination. Species name search included both recognized scientific names and synonyms, placed in parentheses within a same search query (i.e., [“*species name*” OR “*synonym #1*” OR “*synonym #2*” OR…]), to avoid negative effect of taxonomic synonyms on search accuracy (Correia et al., 2018). Searches for the two threats were performed with the commonly used terms in each country (Supplementary material S1). Although the focus of the study were species and threats in the three countries within Europe, Google’s Custom Search Engine API also covered data from their overseas territories. Nevertheless, this is unlikely to substantially affect our results, mainly due to small proportion of the human population of overseas territories (∼4% for France, <1% for United Kingdom).

Data analysis was based on the relative Internet salience of the two threats in relation to each of the species. Relative Internet salience was expressed as the proportion of webpages that include particular species name and the given threat to all webpages that mention that species ([“*species name*” AND “*threat name*”] / [“*species name*”]). Analysis included comparisons of relative Internet salience between the two threats, among national, European and global threatened species pools, between threatened and non-threatened species, among the four species groups, among the three countries, and between species classified as affected by the threat at issue within the IUCN Red List database and those that are not. We also assessed relative Internet salience of the studied species in relation to the two threats, by expressing the Internet salience as the proportion of webpages that includes species name and the given threat to all webpages that mention that threat *([“species n*ame” AN*D “threat n*ame”] / *[“threat n*ame”]).

Since data were not normally distributed (Kolmogorov-Smirnov test, *p*<0.001), statistical analyses were performed using nonparametric tests. Differences between groups were assessed using the Mann-Whitney U test with Bonferroni correction, while the relationship between relative Internet salience and the severity of each threat was assessed using the Spearman’s Rank test.

## 3. Results

Climate change had a significantly higher relative Internet salience than biological invasions. This was consistent among countries and species groups (Figs. 1-4). Furthermore, both threats were better represented among species present in Germany, France, and United Kingdom than among those outside of each of the countries, and they were also more represented among species from Europe than among those distributed outside of Europe (Mann-Whitney U test, *p*<0.01; Fig. 1). Climate change had a significantly higher prominence than biological invasions within amphibians, birds and mammals (Mann-Whitney U test, *p*<0.05), while the differences in the representation of the two threats within reptiles were not significant (Fig. 2). Both threats had significantly higher Internet salience within threatened than within non-threatened species in Germany and United Kingdom (Mann-Whitney U test, *p*<0.01), but not in France (Fig. 3).

**Figure 1.**
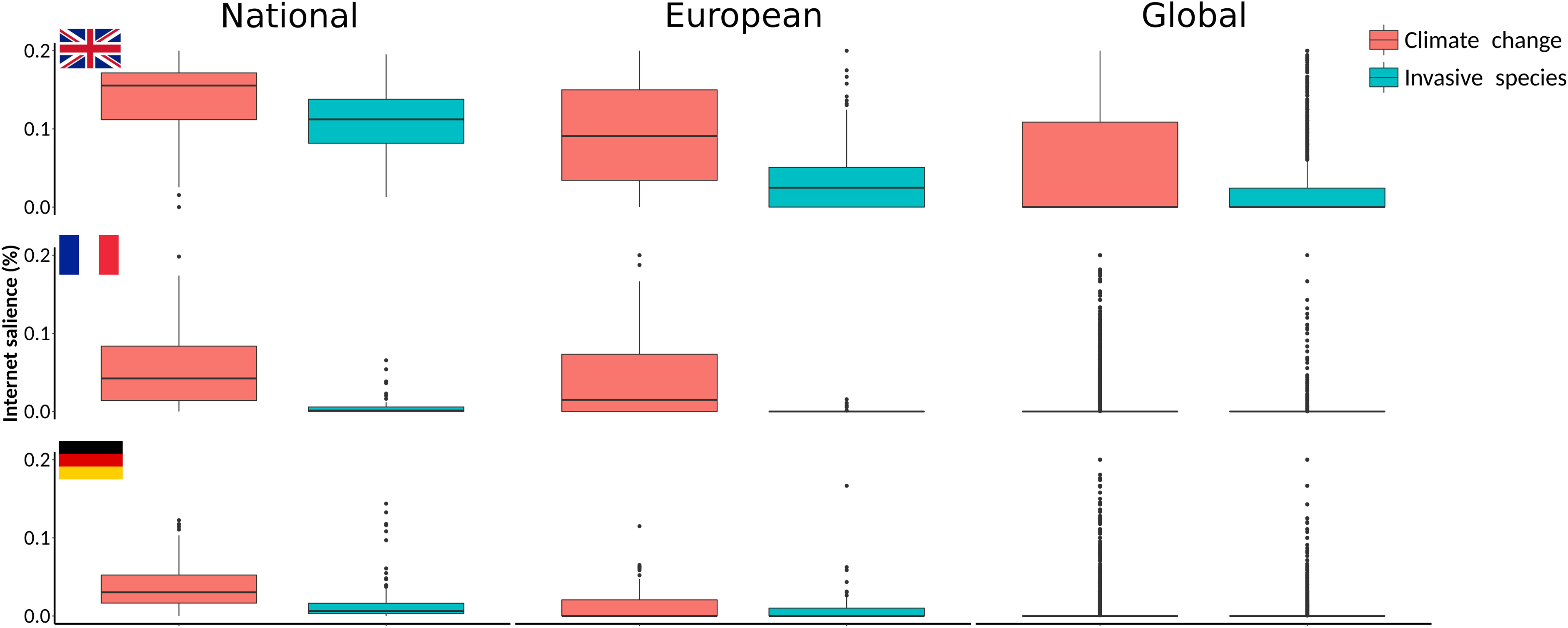
Climate change is a more salient threat on Internet than invasive species; both threats are also better recognized for threatened species within the country than elsewhere in Europe and in the world. Relative Internet salience of climate change and invasive species in relation to threatened species from United Kingdom, France and Germany, as well as on other threatened species of these groups, present elsewhere in Europe and in the world (noted in figure as national, European and global, respectively). Relative Internet salience was expressed as the mean number of webpages retrieved by Internet search within each of the countries for the scientific name of a species and the particular threat, divided by the number of webpages retrieved by searching for the scientific name only.

**Figure 2.**
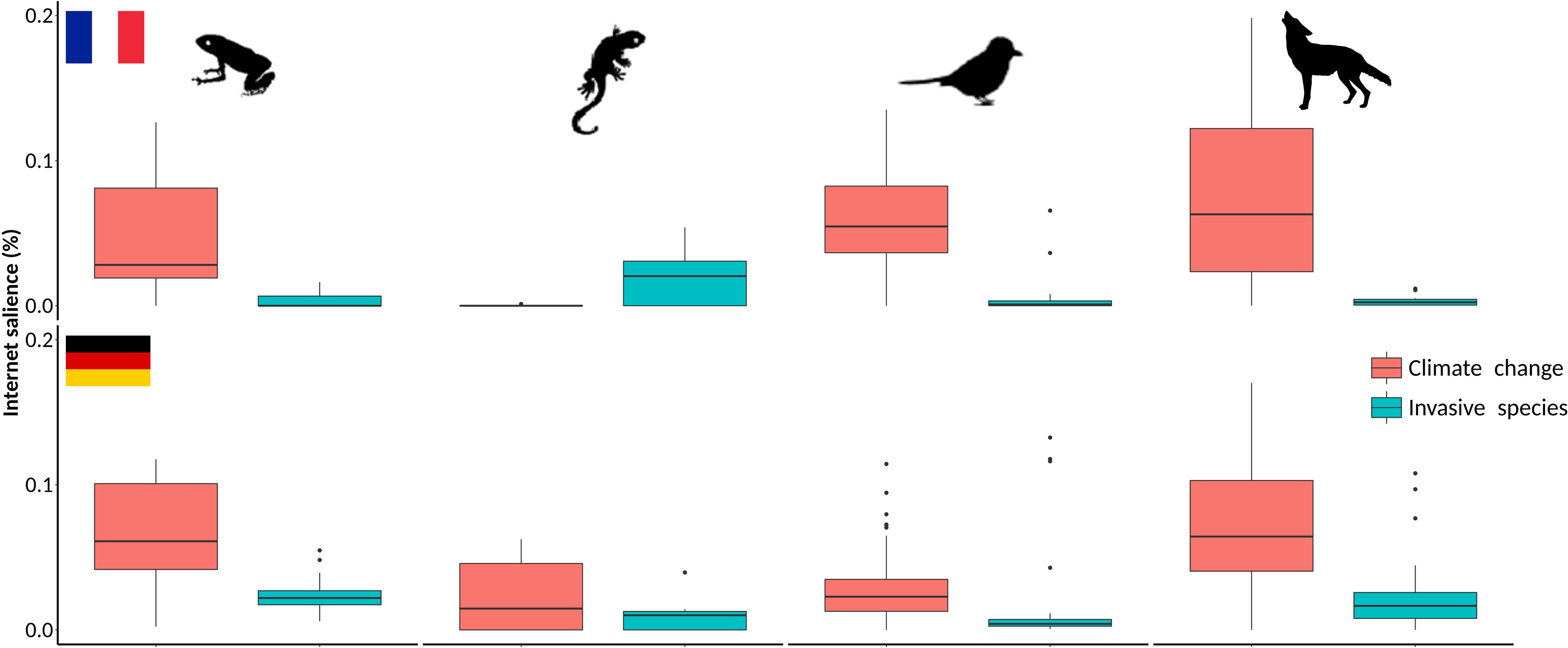
Climate change is better represented on Internet than invasive species in most of the studied species groups and countries. Relative Internet salience of climate change and invasive species in relation to threatened species from the four studied species groups (amphibians, reptiles, birds and mammals) from France and Germany. Relative Internet salience was expressed as the mean number of webpages retrieved by Internet search within each of the countries for the scientific name of a species and the particular threat, divided by the number of webpages retrieved by searching for the scientific name only. Data for United Kingdom were not presented, since the national red list comprised almost exclusively bird species.

**Figure 3.**
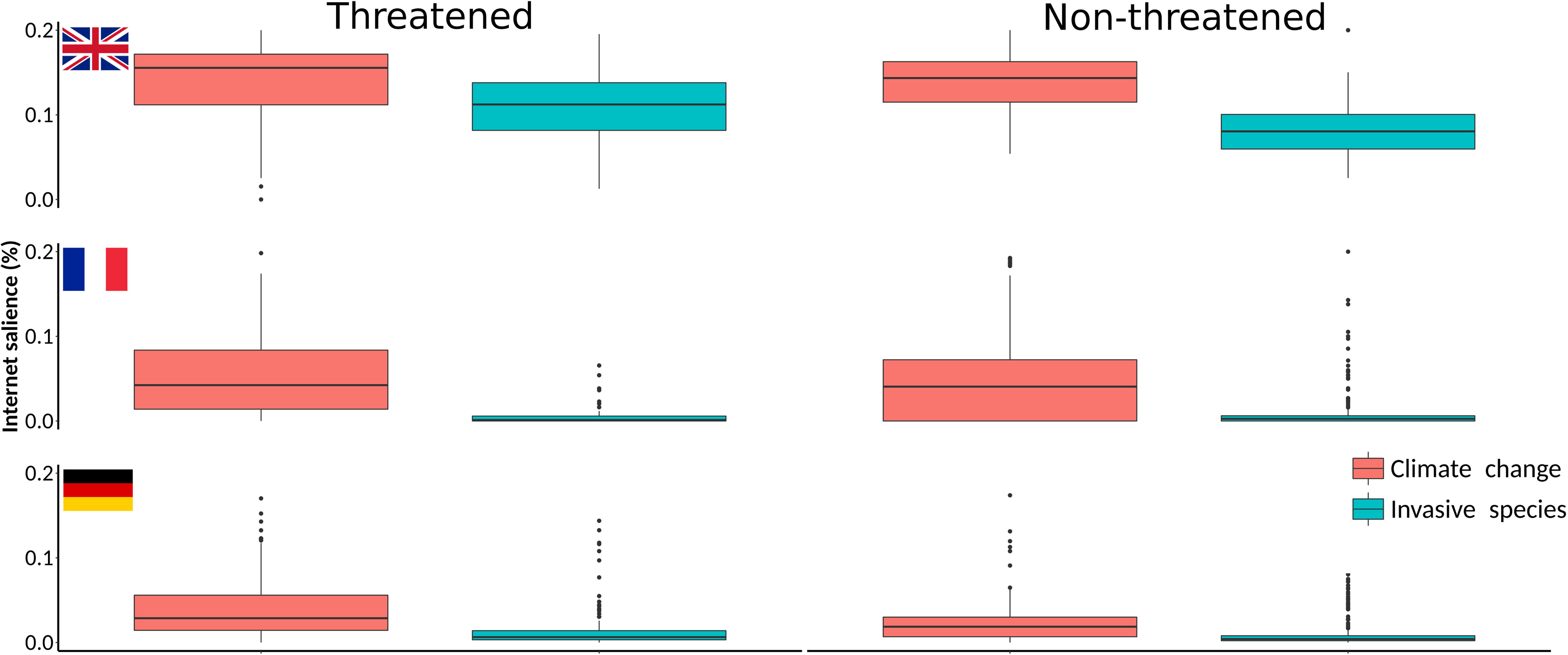
Climate change is a more salient threat on Internet than invasive species, especially among threatened species. Relative Internet salience of climate change and invasive species in relation to threatened and non-threatened species from United Kingdom, France and Germany. Relative Internet salience was expressed as the mean number of webpages retrieved by Internet search within each of the countries for the scientific name of a species and the particular threat, divided by the number of webpages retrieved by searching for the scientific name only.

**Figure 4.**
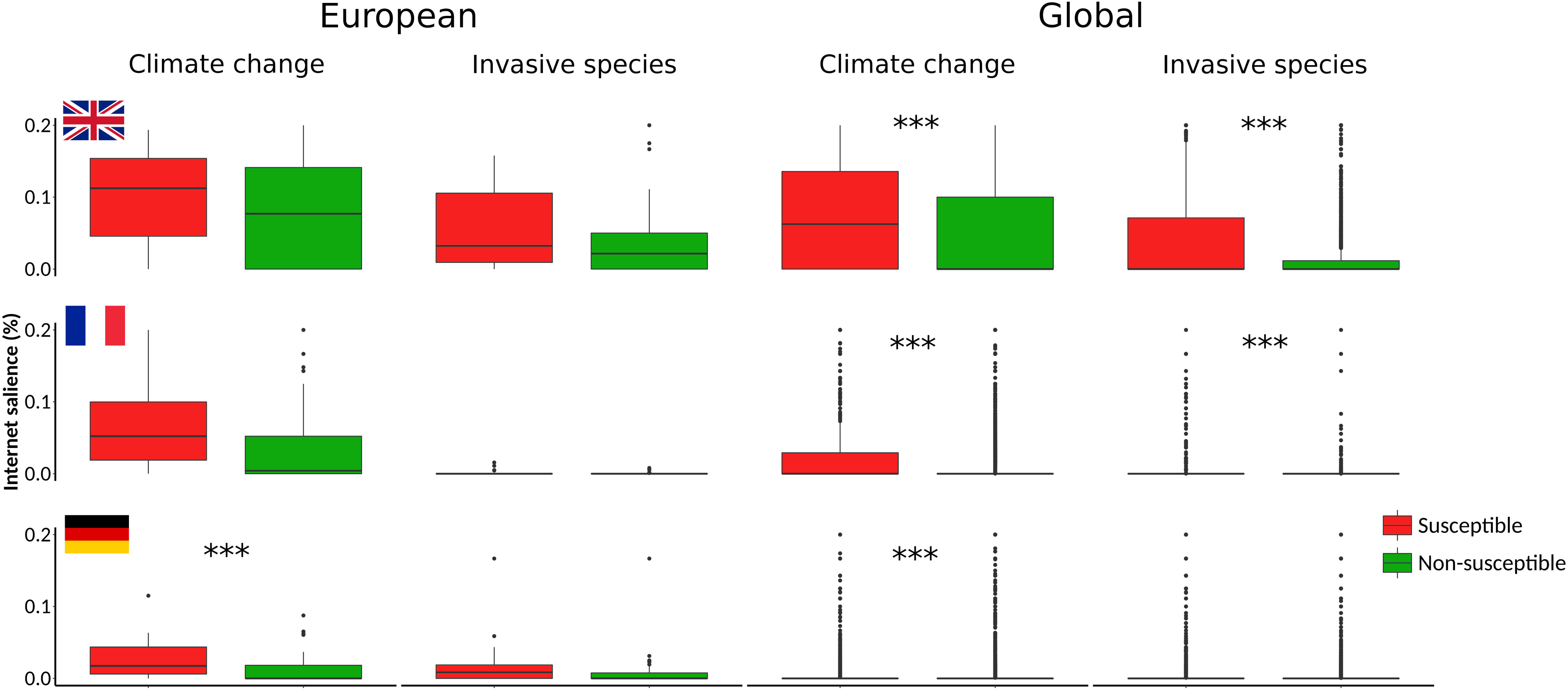
Climate change and invasive species are more salient on Internet for species that are considered susceptible to each of the threats. Relative Internet salience of climate change and invasive species in relation to threatened species within and outside of Europe, which were either classified or not classified as susceptible to the threat at issue within the IUCN Red List database (IUCN, 2017). Relative Internet salience was expressed as the mean number of webpages retrieved by Internet search within each of the three countries for the scientific name of a species and the particular threat, divided by the number of webpages retrieved by searching for the scientific name only. Asterisks indicate significant differences (Mann Whitney U Test with Bonferroni correction, *p* < *α*, where *α* = 0.05 / 12 = 0.00417).

On the other hand, the proportion of pages mentioning a certain threat that also contained names of particular species had the opposite pattern, with biological invasions being better represented than climate change (Table 1). In other words, webpages addressing climate change had less frequent mentions of particular species than those dealing with biological invasions. This was especially prominent for France (Table 1).

**Table 1.** Relationship between the Internet salience of climate change and invasive species when Internet search included the name of one of the threats only, and when it comprised also scientific names of threatened amphibian, reptile, bird and mammal species from United Kingdom, France and Germany.

| Internet search type |  | Internet salience |  | Proportion (%) |
| --- | --- | --- | --- | --- |
|  |  | Threat + scientific name (mean across species) | Threat |  |
| Climate change | UK | 189.1 (0 - 552) | 858,000 | 0.022 (0.000 - 0.064) |
|  | France | 44.3 (0 - 489) | 400,000 | 0.011 (0.000 - 0.122) |
|  | Germany | 49.4 (0 - 723) | 835,000 | 0.006 (0.000 - 0.087) |
| Invasive species | UK | 99.4 (2 - 354) | 192,000 | 0.052 (0.001 - 0.184) |
|  | France | 3.7 (0 - 63) | 1,450 | 0.255 (0.000 - 4.344) |
|  | Germany | 22.4 (0 - 384) | 9,060 | 0.247 (0.000 - 4.238) |

Species that were recognized as susceptible to either climate change or biological invasions within the IUCN Red List database (IUCN, 2017) overall had a better representation of the given threat than those that were not recognized as susceptible. This indicates that the representation of threats on the Internet corresponds well to expert identification. Internet salience of threats was positively, but weakly correlated with threat severity in species impacted by a given threat (Table 2). Such results indicate the potential presence of a pattern, albeit a weak one, where species that are more strongly impacted by either biological invasions or climate change were characterized by a better representation of that threat on the Internet (Table 2).

**Table 2.** Coefficients of correlation (Spearman’s non-parametric correlation test) between the relative Internet salience of climate change and invasive species and the assigned threat severity within the IUCN Red List database (IUCN, 2017). Dataset comprises threatened species from the four studied taxon groups from United Kingdom, France and Germany, classified as susceptible to the threat at issue within this database.

|  | UK | France | Germany |
| --- | --- | --- | --- |
| Climate change | 0.13<sup>**</sup> | 0.12<sup>**</sup> | 0.10<sup>*</sup> |
| Invasive species | 0.16<sup>**</sup> | 0.18<sup>**</sup> | 0.11<sup>**</sup> |

<sup>\*</sup> $p < 0.05$ ; <sup>\*\*</sup> $p < 0.01$

## 4. Discussion

Our results revealed substantial differences in the Internet salience of the two studied threats, with a considerably higher prominence of climate change as a threat in assessed species groups than biological invasions. Such results were expected and intuitive. Although the problem of biological invasions has been known for hundreds of years, the issue of climate change emerged much more recently but quickly gained prominence and has remained the dominant environmental concern over the past few decades (Veríssimo et al., 2014; Capstick et al., 2015). This is also supported by the dominance and an increasing trend in climate change frequency in Internet search queries and media, as opposed to biological invasions and other conservation topics that were characterized by considerably lower and often declining trends (McCallum and Bury, 2013; Nghiem et al., 2016; Legagneux et al., 2018; but see Burivalova et al., 2018; Correia et al., 2019). One of the reasons for a higher visibility of climate change is that it is experienced directly by the public and it has global effects, with easily perceived economic damages, while biological invasions and most of the other threats are perceived as more local, context-dependent and indirect impacts (Legagneux et al. 2018). Research related to climate change is a large, multidisciplinary, field with multiple scientific disciplines and a large number of researchers involved, which consequently translates into more intense and effective science outreach. Visibility of and interest in climate change is also further enhanced by the presence of an intergovernmental panel of scientists on climate change, but probably also by ongoing debates about its causes, effects and very existence. It is also important to note that biological invasions are comparably a less well known and understood process, and consequently less efficiently communicated to the public (Courchamp et al., 2017). On the other hand, activities of the Intergovernmental Science-Policy Platform on Biodiversity and Ecosystem Services (IPBES), including the recent publication of the Global Assessment Report on Biodiversity and Ecosystem Services (IPBES, 2019) and the establishment of a special working group for invasive alien species, are likely to help improve communication about biological invasions. Society needs to be aware and recognize the reality of a certain threat, as well as that it affects particular species, before it can become interested and invested in the related management issues. In this respect, attention toward the impacts of these two threats on different species is also heavily affected by the prominence of climate change skepticism and invasive species denialism (Antilla, 2005; Capstick et al., 2015; Russell and Blackburn, 2016; Ricciardi and Ryan, 2018), as well as by societal perception of particular invasive alien species and their impacts (Shackleton et al., 2018; Garcia-Llorente et al., 2008).

Climate change was however much less associated with species than biological invasions (Table 1), probably due to the way these threats are perceived and addressed in management. For managers in the field, acting directly on climate change is in most cases impossible. Consequently, managers may less frequently mention climate change in actions to be implemented. Climate change is perceived as a broader threat and mostly addressed independently of any species that might be affected. Rare exceptions are species that could be artificially translocated by humans to more suitable climate zones (Hoegh-Guldberg et al., 2008; Thomas, 2011). On the other hand, management of biological invasions often takes into consideration species that are threatened by a particular harmful invader, and such species are usually also explicitly addressed in the management (Jones et al. 2016).

Both threats were characterized by notable differences among countries, species groups and their origin. Either threat had much higher coverage related to species native for a given country than to those from other countries and regions. People are more knowledgeable of their immediate surroundings, and the species familiarity and the range proximity to or overlap with developed nations are recognized drivers of both societal and scientific taxonomic attention (Correia et al., 2016; Jarić et al., 2019a). This is further strengthened by regional differences and gaps in knowledge and information about these threats, due to strong spatial research biases (Bellard and Jeschke, 2016; Bellard et al., 2018). France showed a particular behaviour with invasive species compared to the two other countries, with considerably higher association of particular species with the threat.

Two potentially encouraging results of the study are higher societal interest in the two threats in threatened than within non-threatened species, as well as that the IUCN Red List listing of threats and their severity in different species were reflected in threat coverage among species. This might suggest a good level of societal attention toward threats and proper coverage of scientific work and available knowledge, supporting recent findings that suggest a high level of overlap between scientific and societal taxonomic attentions (Jarić et al., 2019a). However, while knowledge and awareness of threats is a necessary condition for actual conservation engagement, we should bear in mind that it is not always sufficient.

It is important to note some potential limitations of the study, that call for due caution when interpreting results. Use of phrases such as “invasive species” and “climate change” can sometimes be ambiguous and perceived by the public with a different meaning than the one established in science, not necessarily reflecting perception of these threats as conservation issues. For example, people sometimes tend to refer to native species whose populations are on the rise, such as roe deer (*Capreolus capreolus*) or wild boar (*Sus scrofa*), as “invasive species”. However, even though it was not possible to monitor the effects of such ambiguities within the assessed dataset, they are probably too scarce to significantly affect results. Furthermore, the potential problem of global unevenness of the public access and use of Internet was avoided here by focusing on European countries where most of the population are active Internet users. Finally, use of web search engines also requires caution, because the results returned may vary with time, both due to changes in search engine algorithms (e.g. personalization algorithms, webpage indexation) and temporal instability of web pages. Nevertheless, by focusing our analysis on relative differences instead of absolute values should ensure that the overall patterns should be robust to these variations.

One part of our analysis included comparisons of the species vulnerability to threats based on the IUCN Red List threat classification with Internet salience of threats. Reliability of the IUCN Red List threat classification was sometimes questioned, especially regarding climate change threat assignment (Trull et al., 2018). Nevertheless, recent studies have shown good performance of IUCN Red List threat assignment (Keith et al., 2014; Pearson et al., 2014), while a set of objective threat assignment criteria established by IUCN and a wide pool of expert assessors involved in the assessment should ensure that the database represents the best available knowledge (Jarić et al., 2019b).

The observed disparity in the online coverage of climate change and biological invasions supports recent claims that general attention toward biological invasions seems to be limited (Courchamp et al., 2017). In addition, low proportion of Internet salience of climate change and biological invasions when Internet search included scientific names of threatened vertebrates also revealed that species conservation concerns represent small proportion of the overall Internet content. Yet, Internet coverage of the two threats also revealed the presence of two general communication models: climate change is a well-recognized threat, but addressed in a more general way, mostly not associated with specific species; on the other hand, biological invasions are less recognized, but they are more often associated with particular impacted species. There is a need to devise more efficient communication and outreach approaches regarding the threat from biological invasions, especially since our results may imply that a lack of attention is in part the fault of the media. A set of tools from climate change communication that were advocated by Legagneux et al. (2018) for improved communication on biological invasions, including the use of metaphors, iconic species and mobilizing information, could be a step in the right direction.

## Acknowledgements

Authors would like to thank Janet Scott for providing the IUCN Red List data. IJ acknowledges the sponsorship provided by the The J. E. Purkyně Fellowship of the Academy of Sciences of the Czech Republic, Alexander von Humboldt Foundation and the German Federal Ministry of Education and Research (BMBF). RAC is currently supported by a post-doctoral grant from Fundação para a Ciência e Tecnologia (SFRH/BPD/118635/2016) and thanks FCT/MCTES for the financial support to CESAM (UID/AMB/50017/2019), through national funds. FC acknowledges the support by the Invacost research program (ANR and BNP Paribas).

