## Supplementary material S1 for "Societal attention toward extinction threats"

**Supplementary material S1.** Terms used for searches for the two threats for each country.

|  | Climate change | Invasive species |
| --- | --- | --- |
| Germany | ["Klimawandel" OR "Erdewärmung" OR "globale Erwärmung"] | ["invasive Arten"] |
| France | ["changement climatique" OR "dérèglement climatique" OR "réchauffement climatique" OR "réchauffement planétaire" OR "réchauffement de la planète" OR "réchauffement de la terre"] | ["espèces envahissantes" OR "espèces invasives"] |
| United Kingdom | ["climate change" OR "global warming"] | ["invasive species"] |
